# Copy-number signatures and mutational processes in ovarian carcinoma

**DOI:** 10.1101/174201

**Authors:** Geoff Macintyre, Teodora E. Goranova, Dilrini De Silva, Darren Ennis, Anna M. Piskorz, Matthew Eldridge, Daoud Sie, Liz-Anne Lewsley, Aishah Hanif, Cheryl Wilson, Suzanne Dowson, Rosalind M. Glasspool, Michelle Lockley, Elly Brockbank, Ana Montes, Axel Walther, Sudha Sundar, Richard Edmondson, Geoff D. Hall, Andrew Clamp, Charlie Gourley, Marcia Hall, Christina Fotopoulou, Hani Gabra, James Paul, Anna Supernat, David Millan, Aoisha Hoyle, Gareth Bryson, Craig Nourse, Laura Mincarelli, Luis Navarro Sanchez, Bauke Ylstra, Mercedes Jimenez-Linan, Luiza Moore, Oliver Hofmann, Florian Markowetz, Iain A. McNeish, James D. Brenton

**Author notes:** These authors contributed equally to this work (Geoff Macintyre, Teodora E. Goranova). Co-corresponding authors: Florian Markowetz, Iain McNeish. James Brenton.

## Abstract

Genomic complexity from profound copy-number aberration has prevented effective molecular stratification of ovarian and other cancers. Here we present a method for copy-number signature identification that decodes this complexity. We derived eight signatures using 117 shallow whole-genome sequenced high-grade serous ovarian cancer cases, which were validated on a further 497 cases. Mutational processes underlying the copy-number signatures were identified, including breakage-fusion-bridge cycles, homologous recombination deficiency and whole-genome duplication. We show that most tumours are heterogeneous and harbour multiple signature exposures. We also demonstrate that copy number signatures predict overall survival and changes in signature exposure observed in response to chemotherapy suggest potential treatment strategies.

---

The discrete mutational processes that drive copy-number change in human cancers are not readily identifiable from genome-wide sequence data. This presents a major challenge for the development of precision medicine for cancers that are strongly dominated by copy-number changes, including high-grade serous ovarian (HGSOC), oesophageal, non-small-cell lung and triple negative breast cancers^1^. These tumours have low frequency of recurrent oncogenic mutations, few recurrent copy number alterations and highly complex genomic profiles^2^.

HGSOCs are poor prognosis carcinomas with ubiquitous *TP53* mutation^3^. Despite efforts to discover new molecular subtypes and targeted therapies, overall survival has not improved over two decades^4^. Current genomic stratification is limited to defining homologous recombination-deficient (HRD) tumours^5–7^, and classification using gene expression does not currently have clinical utility^8,9^. Detailed genomic analysis using whole genome sequencing has shown frequent loss of *RB1, NF1* and *PTEN* by gene breakage events^10^ and enrichment of amplification associated fold-back inversions in non-HRD tumours^11^. However, none of these approaches has provided a broad mechanistic understanding of HGSOC, reflecting the challenges of detecting classifiers in extreme genomic complexity.

Recent algorithmic advances have enabled interpretation of complex genomic changes by identifying mutational signatures - genomic patterns that are the imprint of mutagenic processes accumulated over the lifetime of a cancer cell^12^. For example, UV exposure or mismatch repair defects induce distinct single nucleotide variant (SNV) signatures^12^, whilst signatures encoded by structural variants (SVs) can summarise different types of HRD^13^. Importantly, these studies show that tumours typically harbour multiple mutational processes requiring computational approaches that can robustly identify coexistent mutational signatures. Quantification of the exposure of a tumour to specific mutational signatures provides a rational framework to personalise therapy^14^ but currently is not readily applicable to copy-number driven tumours. We hypothesized that specific features of copy number abnormalities could also reflect the imprint of mutational processes on the genome, and we thus developed methods to identify copy-number signatures using shallow whole genome sequencing for disease stratification in HGSOC.

## Results

### Identification and validation of copy-number signatures

To identify copy-number (CN) signatures, we sequenced 300 primary and relapsed HGSOC samples from 142 patients in the BriTROC-1 cohort^15^ (Supplementary Figure 1). We combined low-cost shallow whole-genome sequencing (sWGS; 0.1×) with targeted amplicon sequencing of *TP53* to generate absolute copy-number profiles. Copy-number profiles from 117 patients were used for copy-number signature identification (Figure 1).

**Figure 1.**
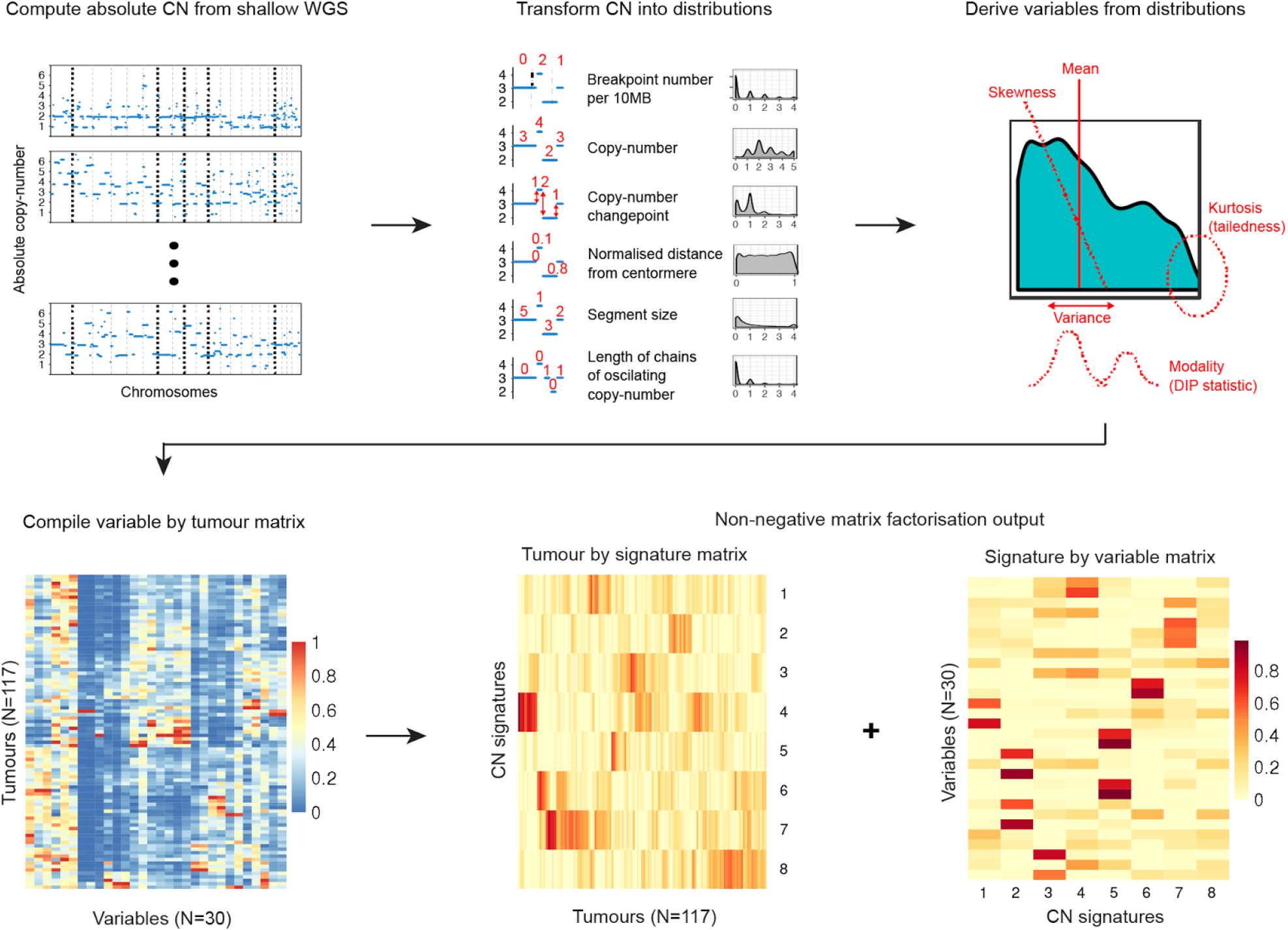
Copy-number signature identification from shallow whole genome sequence data. Step 1: Absolute copy-numbers are derived from sWGS data; Step 2: genome-wide distributions of six fundamental copy-number features are computed; Step 3: the shape of each distribution is computed using the mean, variance, skewness, kurtosis, and modality; Step 4: the data are represented as a matrix with 30 (=6x5) variables per tumour. Step 5: Non-negative matrix factorisation is applied to variable-by-tumour matrix to derive the tumour-by-signature matrix and the signature-by-variable matrix.

For each sample, we computed the genome-wide distributions of six fundamental CN features: the number of breakpoints per 10MB, the copy-number of segments, the size of segments, the difference in CN between adjacent segments, the distance of breakpoints from the centromere, and the lengths of oscillating CN segment chains. These features were selected as hallmarks of previously reported genomic aberrations, including breakage-fusion-bridge cycles^16^, chromothripsis^17^ and tandem duplication^18,19^. We summarised each feature using measures of the shape of its distribution: mean, variance, skewness (asymmetry), kurtosis (weight of tails) and modality (number of peaks), thus describing each tumour sample with 30 variables (Figure 1).

To identify copy-number signatures from these data, we used non-negative matrix factorisation (NMF)^20^, a method previously used for analysing SNV signatures^12^. NMF identified eight CN signatures (Figure 1), as well as their defining features and their exposures in each sample. The optimal number of signatures was chosen using a consensus from 1000 initialisations of the algorithm and 1000 random permutations of the data combining four model selection measures (Supplementary Figure 2). We found highly similar variable weights for the signatures in two independent cohorts of 95 whole-genome sequenced HGSOC samples from ICGC^21^ and 402 SNP array profiled HGSOC samples^22^, demonstrating the robustness of both the methodology and the copy number features across different sample sets (Supplementary Figure 3, P<0.005, median r^2^=0.85. Supplementary Table 1).

### Linking copy-number signatures with underlying mutational processes

Previous studies in HGSOC have identified dominant genomic patterns such as tandem-duplication^19^ or genomic instability^10,23^. Inspection of WGS data from 8 cases with a predominant copy-number signature showed that CN signatures 1, 7 and 8 were characterised by high genome complexity (Supplementary Figure 4), with signature 7 suggestive of a tandem duplicator phenotype^19^.

However, the vast majority of cases exhibited multiple signature exposures suggesting that the complex genotypes in HGSOC are shaped by several mutational processes (see below). Use of NMF was able to reduce this genomic complexity into its constituent components, allowing the linkage of individual copy-number signatures to mutational processes. Specifically, the weights identified by NMF were used to determine which pattern of global or local copy-number change defined each signature. For example, for CN signature 3, the highest weights were observed for the skewness and kurtosis variables for the CN feature distribution describing breakpoint distance from the centromere (Figure 2a). This suggested frequent breakpoints close to the end of the chromosomes, which are characteristic of peri-telomeric breaks from breakage-fusion-bridge (BFB) events^16^. To test this hypothesis, we correlated CN signature 3 exposures with other genomic features from deep WGS (Figure 2b, Supplementary Figures 5 and 6). CN signature 3 was correlated with sequencing estimates of telomere length (r^2^=-0.21, P=0.08, multiple testing corrected) and positively correlated with age-associated SNV signature 1 (r^2^=0.35, P=0.0004), consistent with BFB events. In addition, CN signature 3 was positively correlated with amplification-associated fold-back inversion structural variants (r^2^=0.34, P=0.07), which have been strongly implicated in BFB events^24^ and have been associated with inferior survival in HGSOC^11^. Taken together, these data provide independent evidence for BFB as the underlying mechanism for CN signature 3.

**Figure 2.**
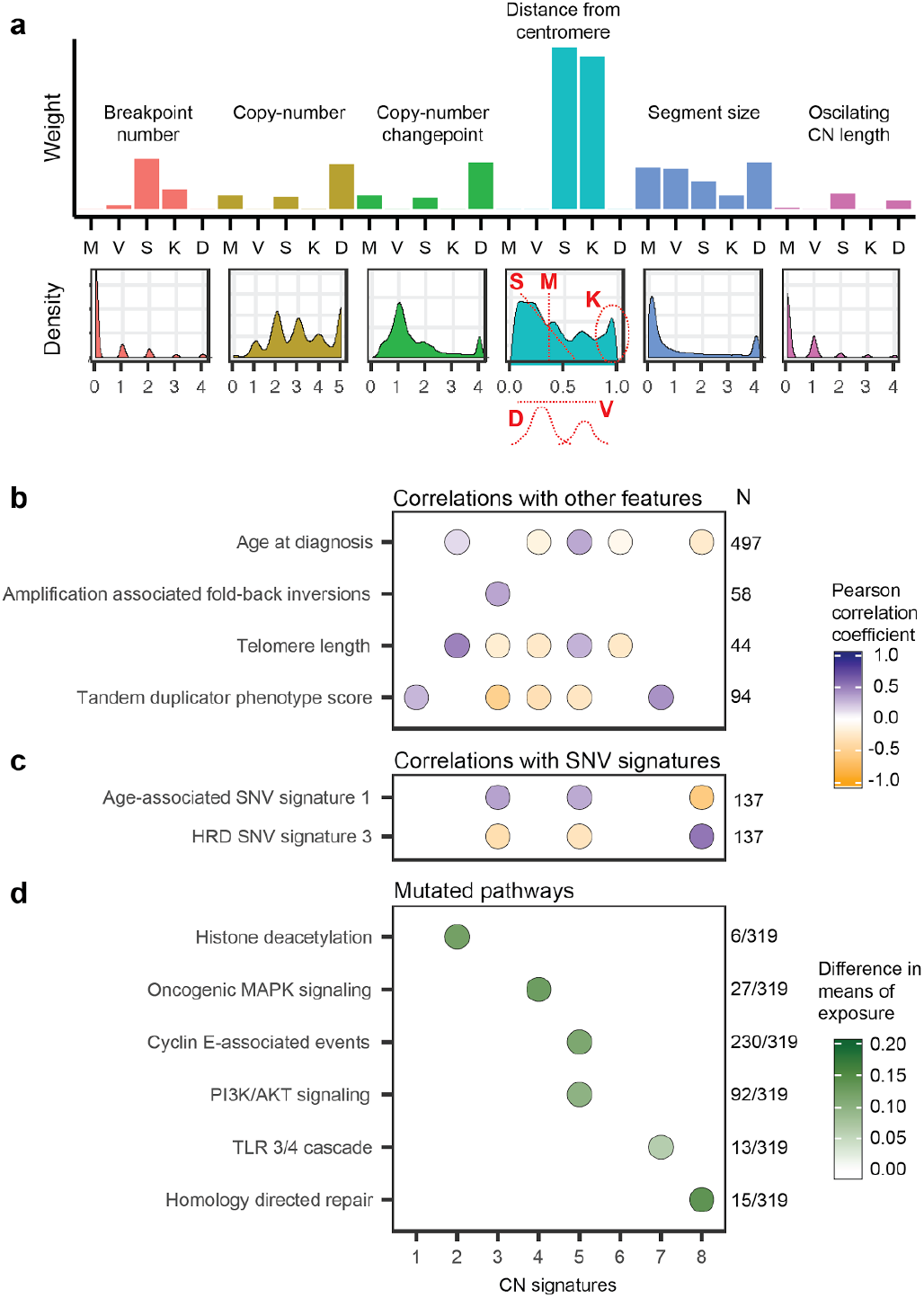
Linking copy-number signatures with mutational processes. **a** Variable weights for copy number signature 3. Barplots (upper panel) are grouped by copy number feature and show weights for each variable. Lower panel shows genome-wide distribution (density) of each copy number feature across all signatures. The distribution for the feature distance from centromere (turquoise) is annotated to show the variables summarizing the distribution (M, Mean; V, Variance; S, Skew; K, Kurtosis; D, DIP statistic). Plots for all signatures are shown in Figure 3 and Supplementary Figure 7. **b** Associations between signature exposures and other genomic features. Purple indicates positive correlation and orange negative correlation. **c** Associations between signature exposures and SNV signature. Purple indicates positive correlation and orange negative correlation. **d** Different mean signature exposures in reactome pathways between mutant and non-mutant cases (FDR P<0.1, one-sided t-test). Colour scale indicates extent of difference. Numbers at the right of each panel indicate cases included in each analysis. Dots only show associations with significant correlation coefficient (FDR P<0.1).

We systematically applied the same approach to the remaining signatures to identify other statistically significant genomic associations using a false discovery rate <0.1 (Figure 2b, Figure 3, Supplementary Figures 5, 6 and 7). For two CN signatures, 1 and 6, we did not find evidence for specific underlying mechanisms. For three signatures, we identified significant associations with signalling pathways implicated in DNA damage. CN signature 2 was associated with mutations in genes involved in histone deacetylation (P=0.007, one-sided t-test) and lengthening of telomeres (r^2^=0.47, P=3e-05); histone deacetylase (HDAC) activity is closely linked to DNA damage responses^25^, specifically via ATM regulation^26^, whilst inhibition of HDACs can induce profound DNA damage^27^ with upregulation of error-prone non-homologous end-joining^28^. CN signature 4 was enriched in cases with oncogenic MAPK signalling including *NF1* loss and mutated *KRAS* (P=0.003, one-sided t-test). MAPK signalling has previously been shown to induce chromosomal instability as a result of aberrant G2 and mitotic checkpoint controls and missegregation^29,30^. CN signature 7 was enriched in patients with high tandem duplicator phenotype scores (r^2^=0.42, P=1e-04) and mutations in the Toll-like receptor (TLR) pathway (P=0.06, one-sided t-test). TLR4 activity regulates expression of Ku70^31^, whilst persistent TLR signalling leads to mitotic defects and potent DNA damage responses^32^, suggesting possible links between TLR pathway mutations and CN abnormalities.

**Figure 3.**
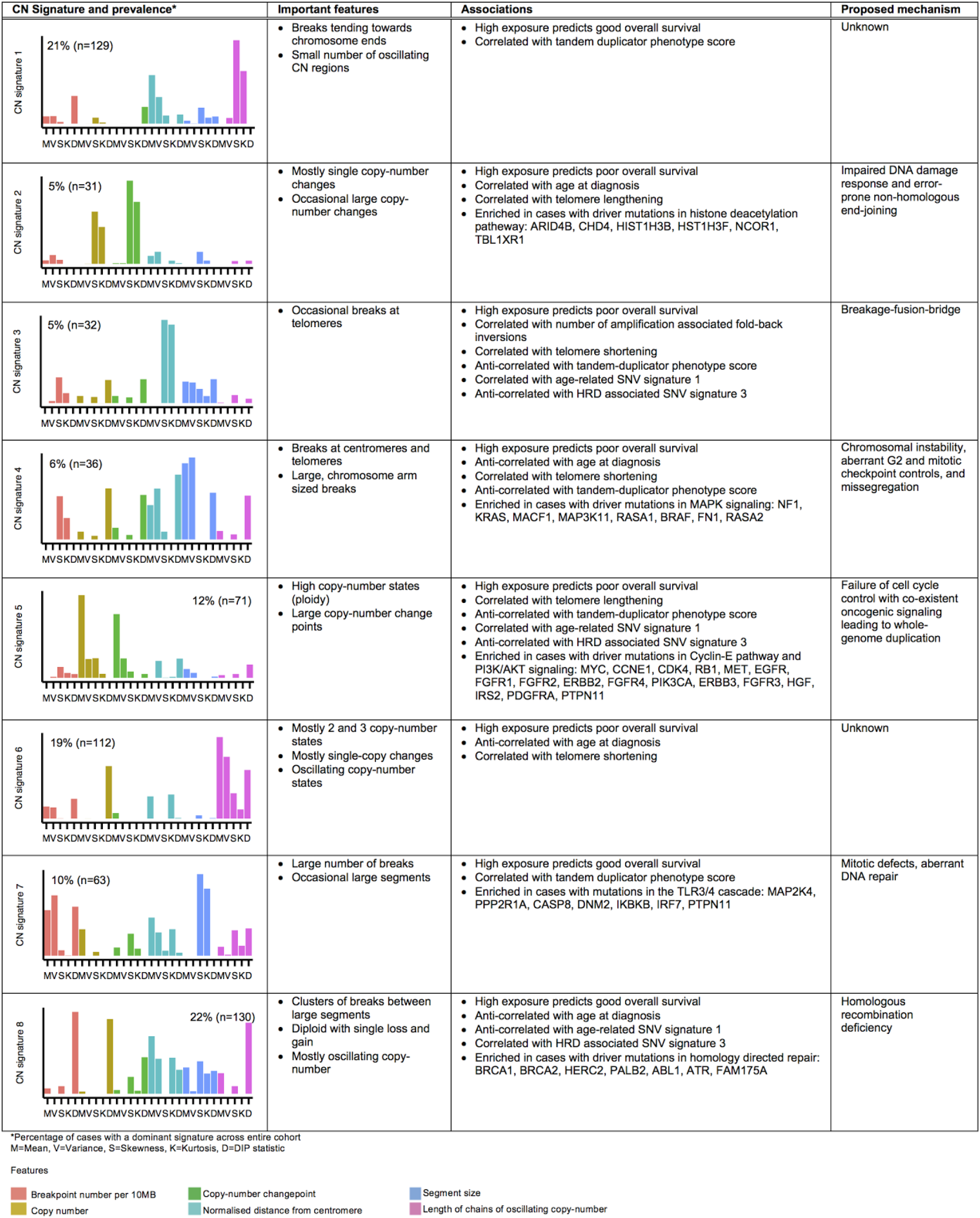
Copy-number signatures in HGSOC.

We found the strongest evidence for underlying mutational processes for signatures 5 and 8. CN signature 5 was found at significantly higher exposures (P=1.7e-16, one-sided t-test) in tumours with aberrant cell cycle control, including either amplification of *CCNE1, CCND1, CDK4* or *MYC*, or deletion/inactivation of *RB1*. In addition, signature 5 was enriched in tumours with activated PI3K/AKT signalling (P=1.9e-07, one-sided t-test) through mutation of *PIK3CA* or amplification of *EGFR, MET, FGFR3* and *ERBB2*, suggesting that the underlying mechanism for this signature is failure of cell cycle control with co-existent oncogenic signalling. Exposure to CN signature 5 was positively correlated with age at diagnosis (r^2^=0.33, P=3e-13) and age-related SNV signature 1^12^ (r^2^=0.33, P=5e-04). CN signature 5 was characterised by large values in the copy-number change-point distribution, suggesting that it represents a single, late whole-genome duplication event^33^.

CN signatures 5 and 8 also showed mutually exclusive associations (Figure 2b and 2c). Signature 8 had significantly higher exposures in cases with mutations in *BRCA1/2* or other homologous recombination (HR) pathway genes (P=1.4e-05, one-sided t-test) and was positively correlated with SNV signature 3 (associated with *BRCA1/2* mutations, r^2^=0.53, P=4e-10). It was also negatively correlated both with age at diagnosis (r^2^=-0.23, P=8e-07) and the age-related SNV signature 1 (r^2^=-0.57, P=7e-12), suggesting early age of disease onset. By contrast, signature 5 was negatively correlated with SNV signature 3 and positively correlated with age-related SNV signature 1. Together, these results are consistent with HR deficiency being the underlying mechanism for signature 8, and confirm previous data showing mutual exclusivity of *CCNE1* amplification and defective HR^34^.

### Copy-number signatures predict overall survival

Using a combined dataset of 545 samples with full clinical annotation, we explored the association between individual signature exposures and overall survival (Figure 4a). In a multivariate Cox proportional hazards model trained on 347 cases and tested on the remaining 198, CN signatures 1, 7 and 8 had a positive influence on survival, while the remaining five CN signatures had a negative influence (Training: P=0.005, log-rank test; stratified by age and cohort; Test: P=0.03, C-index=0.57, 95% Cl:0.51-0.62). Although there was a high frequency of cases with multiple signature exposures, stratifying patients by their most dominant signature exposure showed significant survival differences (Figure 4b; Training: P=7.6e-5, log-rank test; Test: P=1.1e-4, log-rank test).

**Figure 4.**
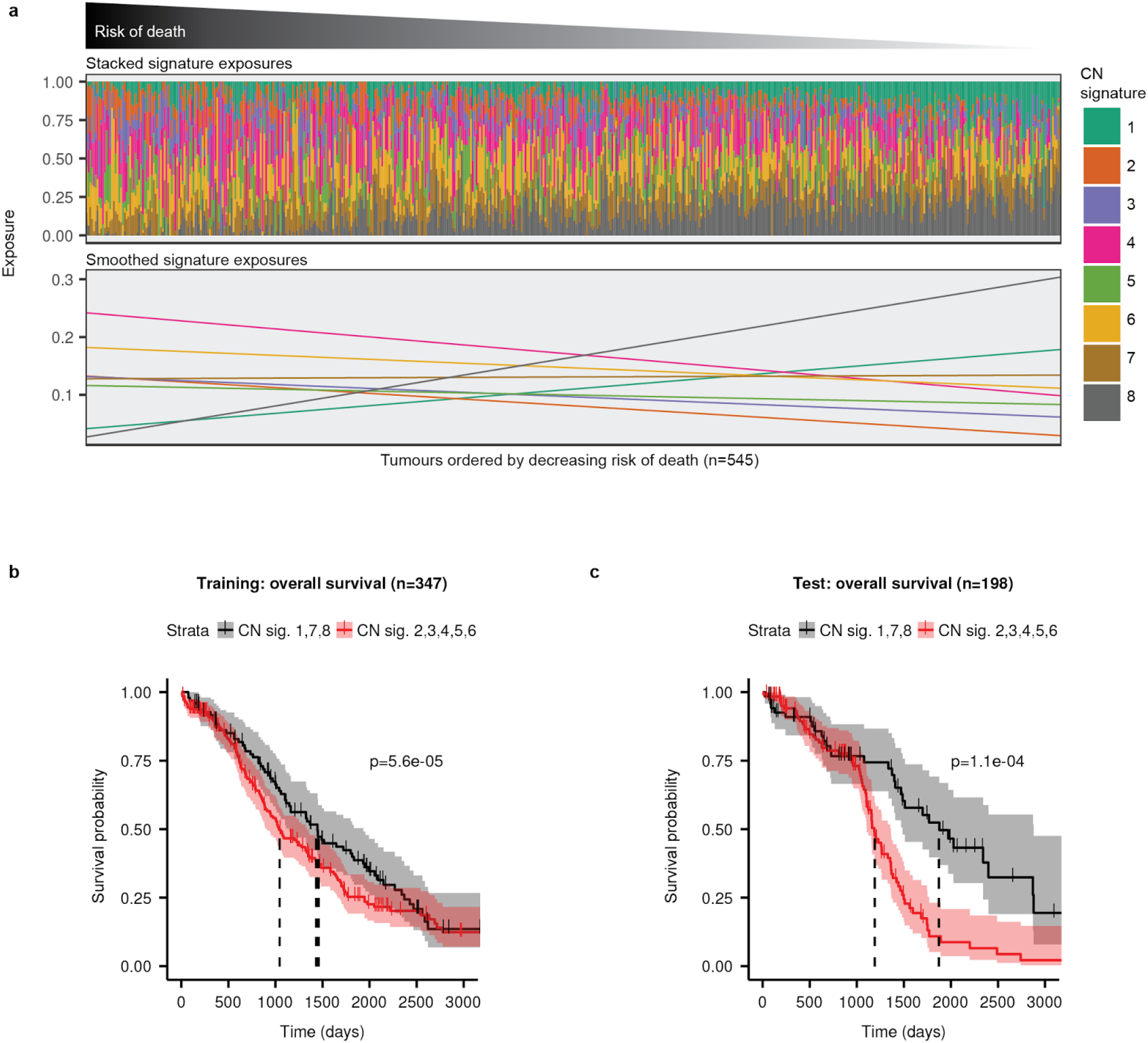
Association of survival with copy-number signatures. **a** Stacked barplots in upper panel show CN signature exposures for each patient. Patients were ranked by risk of death estimated by a multivariate Cox proportional hazards model stratified by age and cohort, with CN signature exposures as covariates. The smoothed exposures for each CN signature are shown in the lower panel. **b** and **c** Survival analysis of patients stratified by dominant signature exposure. Kaplan-Meier curves of overall survival probabilities for patients in b training cohort and c test cohort. Each cohort was divided into two groups based on maximum observed signature exposure (Red: Signatures 2, 3, 4, 5, or 6; Black: Signatures 1, 7 or 8). Black dotted lines indicate the median survival for each group. The p-values reported are from a log-rank test. Shaded areas represent 95% confidence intervals.

### Copy-number signatures indicate response to chemotherapy

We next analysed temporal and treatment effects on copy-number signature exposure by using diagnosis and relapse samples from the BriTROC-1 study (Figure 5a, 152 samples from 116 patients, including 36 with matched sample pairs)^15^.

**Figure 5.**
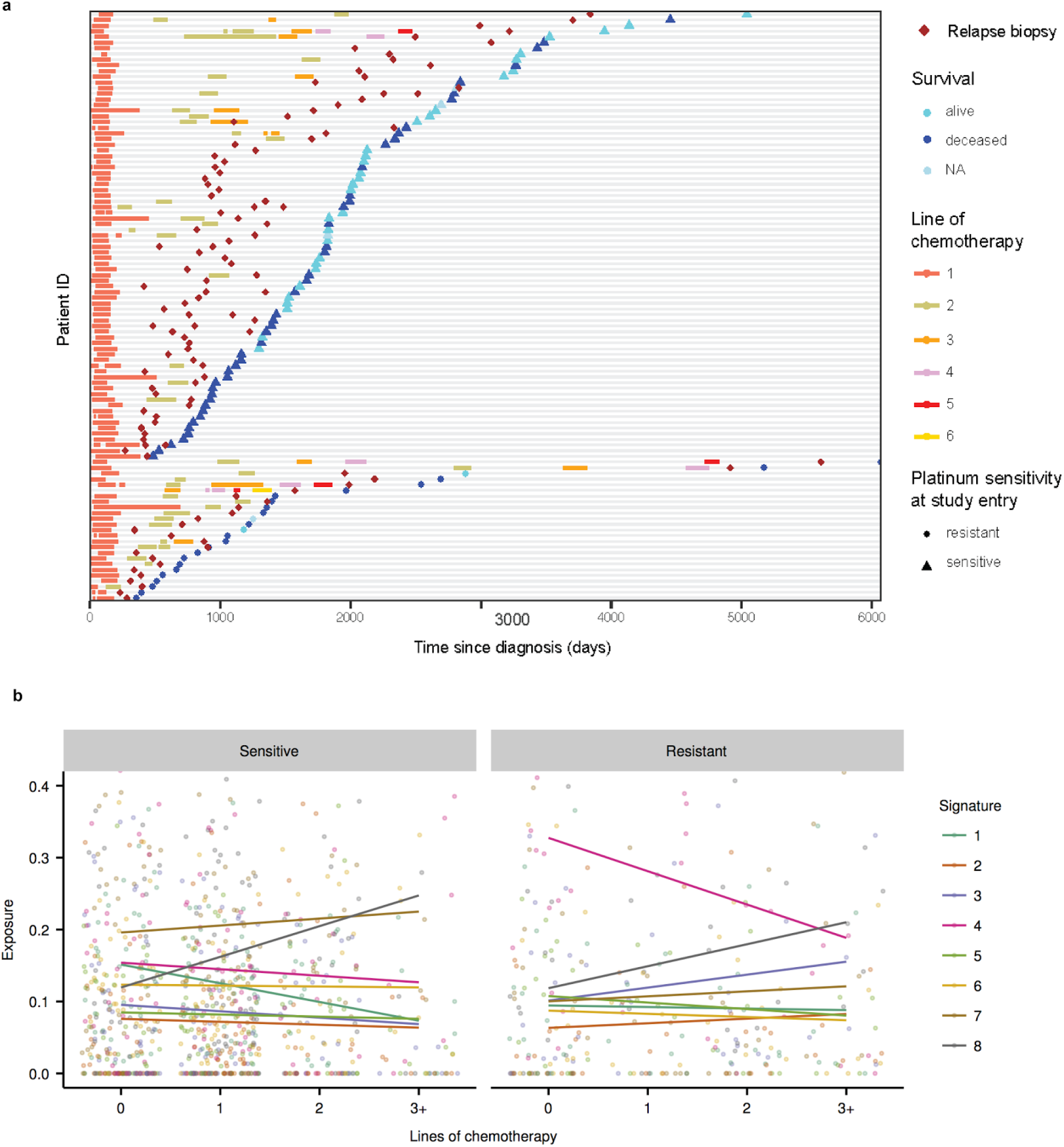
Changes in copy-number signature exposures during treatment. **a** Dot plot of treatment periods and overall survival for 116 BriTROC-1 patients, ranked by overall survival and platinum-sensitive and platinum-resistant relapse. At study entry, patients were classified as having either platinum-sensitive relapse or platinum-resistant relapse based on their preceding therapy. Start and stop dates for line 1 of chemotherapy were missing for three patients. **b** Changes in copy-number signatures across multiple lines of chemotherapy. Points represent signature exposures for BriTROC-1 samples (152 samples from 116 patients) taken at diagnosis (0 prior lines of chemotherapy) and at point of study entry according to number of prior lines of chemotherapy. Coloured lines show the linear model fit for each signature. Samples have been divided into platinum-sensitive (N=117) and platinum-resistant relapse (N=35) samples.

Platinum-resistant tumours showed significantly higher exposures to copy-number signature 4 at diagnosis compared to platinum-sensitive tumours (Figure 5b, Supplementary Figure 8) (P=0.01, one-sided t-test). This suggests that copy-number signature 4 is predictive of early, platinum-resistant relapse. Furthermore, its apparent decrease over multiple lines of therapy reflects the poor survival of patients with high levels of this signature (Figure 4a).

Platinum-sensitive tumours showed consistently high exposure levels to signature 7 across multiple lines of therapy, suggesting that it is broadly associated with good prognosis and response to chemotherapy (Figure 5b). In addition, sensitive tumours that survived multiple (3+) lines of therapy showed enrichment of copy-number signature 8 (associated with HRD), an observation that was also recapitulated in the 36 patients with paired diagnosis and relapse samples (P=9.2e-5, two-sided paired t-test, Supplementary Figure 8).

## Discussion

Copy-number signatures provide a framework that is able to rederive the major defining elements of HGSOC genomes, including defective HR^22^, amplification of cyclin E^8^, and amplification-associated fold-back inversions^11^. In addition, the CN signatures show significant associations with known driver gene mutations in HGSOC and provide increased statistical power to detect novel associations with gene mutations. We derived signatures using inexpensive shallow whole genome sequencing of DNA from core biopsies, providing a rapid path to clinical implementation. Copy-number signatures open new avenues for clinical trial design by allowing stratification of patients based upon contributions from underlying mutational processes including breakage-fusion-bridge cycles and oncogenic NF1/KRAS and PI3K/AKT signalling.

We found that almost all patients with HGSOC demonstrated a mixture of signatures indicative of combinations of mutational processes. These results suggest that early *TP53* mutation, the ubiquitous initiating event in HGSOC, may permit multiple mutational processes to co-evolve, potentially simultaneously. Even in the context of an additional early driver event such as *BRCA2* mutation in germline carriers, a diverse and variable number of CN signatures is possible (Supplementary Figure 9). Such exposure to multiple signatures may alter the risk of developing therapeutic resistance to agents that target a single process such as HRD.

High exposure to signature 4 at diagnosis, which is characterised by oncogenic MAPK signalling (including *NF1, KRAS* and *NRAS* mutation), predicts subsequent platinum-resistant relapse. Indeed, three of the seven matched platinum-resistant sample pairs came from patients who relapsed after only one prior line of therapy, implying the presence of powerful intrinsic resistance mechanisms that can be readily identified at the time of diagnosis. In addition, we have previously demonstrated the expansion of a resistant subclonal Λ/F7-deleted population following chemotherapy treatment in HGSOC^35^, and it is intriguing that there is marked exposure to CN signature 4 in germline *BRCA2* mutated cases (Supplementary Figure 9). Thus, MAPK signalling may be an important target, especially in first line treatment, to prevent emergence of platinum-resistant disease.

Marked enrichment in copy-number signature 8 exposure, associated with HRD, was observed with increasing lines of chemotherapy, especially in those with platinum-sensitive relapse. These results confirm prior data showing that *BRCA1/2* mutation is associated with both long survival in HGSOC^36^ and extended responses to PARP inhibition^37^. The enrichment in signature 8 may be explained by therapy-induced selection of clones with defective HR or therapy-induced rearrangements that amplify the signature of HRD, as well as long-term survival of patients with exclusive HRD. In the ARIEL2 study^7^, where genomic loss-of-heterozygosity (LOH) was used to predict HRD, 14.5% of cases classified as LOH-low at diagnosis were found to be LOH-high at relapse, suggesting some degree of acquired chemotherapy-induced genomic damage and/or clonal selection in these cases. Nonetheless, a high exposure to CN signature 8 may indicate tumours with the potential to respond multiple times to platinum or PARP inhibitor therapy.

Future studies on larger sample collections are needed to refine CN signature definitions and interpretation. The application of our approach to other tumour types is likely to extend the set of signatures beyond the robust core set identified here. Basal-like breast cancers and lung squamous cell carcinoma, which also have high rates of *TP53* mutation and genomic instability^2^, are promising next targets. Other limitations are technical: we integrated data from three sources, using three different pre-processing pipelines, and the ploidy determined by different pipelines can have a significant effect on the derived signatures. For example, high-ploidy CN signature 5 was predominantly found in the sequenced samples that underwent careful manual curation to identify whole-genome duplication events. When extending to larger sample sets, a unified processing strategy with correct ploidy determination is likely to produce improved signature definitions.

In summary, our results represent a substantial advance in deconvoluting the profound genomic complexity of HGSOC, allowing robust molecular classification of a disease that lacks classic driver oncogene mutations or recurrent copy-number changes. By dissecting the mutational forces shaping highly aberrant copy-number, our study paves the way to understanding and categorising extreme genomic complexity, as well as revealing the evolution of tumours as they relapse and acquire resistance to chemotherapy.

## Author contributions

Conceptualisation: GM, TEG, FM, IMcN, JDB; Study conduct: SD, RMG, ML, EB, AM, AW, SS, RE, GDH, AC, CG, MH, CF, HG, DM, AHo, GB, IMcN, JDB; Investigation: TEG, DE, AMP, LAL, AHa, CW, CN, LMi, LNS, MJL, LMo, AS, JP; Formal analysis: GM, TEG, DDS, ME, DS, BY, OH, FM; Methodology and software: GM, DDS, FM; Writing: GM, TEG, DDS, FM, IMcN, JDB

## Acknowledgements

The BriTROC-1 study was funded by Ovarian Cancer Action (to IMcN and JDB, grant number 006). We would like to acknowledge funding and support from Cancer Research UK (grant numbers A15973, A15601, A18072, A17197 and A19274), the Universities of Cambridge and Glasgow, National Institute for Health Research Cambridge Biomedical Research Centre, National Cancer Research Network, the Experimental Cancer Medicine Centres at participating sites, the Beatson Endowment Fund and Hutchison Whampoa Limited. The funders had no role in study design, data collection and analysis, decision to publish or preparation of the manuscript. We thank the Biorepository, Bioinformatics, Histopathology and Genomics Core Facilities of the Cancer Research UK Cambridge Institute and the Pathology Core at the Cancer Research UK Beatson Institute for technical support. We would like to thank members of PCAWG Evolution and Heterogeneity Working Group for the consensus copy-number analysis, PCAWG Structural Variation Working Group for the consensus structural variants and PCAWG Technical Working Group for annotating driver mutations in the 95 ICGC samples.

## Online Methods

### Patients and samples

The BriTROC-1 study has been described previously^15^. The study is sponsored by NHS Greater Glasgow and Clyde and ethics/IRB approval was given by Cambridge Central Research Ethics Committee (Reference 12/EE/0349). The study enrolled patients with recurrent ovarian high-grade serous or grade 3 endometrioid carcinoma who had relapsed following at least one line of platinum-based chemotherapy and whose disease was amenable either to image-guided biopsy or secondary debulking surgery. At study entry, patients were classified as having either platinum-sensitive relapse (i.e. relapse six months or more following last platinum chemotherapy) or platinum-resistant relapse (i.e. relapse less than six months following prior platinum chemotherapy). All patients provided written informed consent. Access to archival diagnostic formalin-fixed tumour was also required. Survival was calculated from the date of enrolment to the date of death or the last clinical assessment, with data cutoff at 1 December 2016. At subsequent relapse or progression after chemotherapy following study entry, patients could optionally have a second biopsy under separate consent.

### Tagged-amplicon sequencing

DNA was extracted from relapsed biopsies and archival diagnostic tissue, and mutation screening of *TP53, PTEN, EGFR, PIK3CA, KRAS* and *BRAF* was performed using tagged-amplicon sequencing as previously described^15^.

### Shallow whole genome sequencing (sWGS)

Libraries for sWGS were prepared from 100ng DNA using modified TruSeq Nano DNA LT Sample Prep Kit (Illumina) protocol^38^. Quality and quantity of the libraries were assessed with DNA-7500 kit on 2100 Bioanalyzer (Agilent Technologies) and with Kapa Library Quantification kit (Kapa Biosystems) using to the manufacturer’s protocols. Sixteen to twenty barcoded libraries were pooled together in equimolar amounts and each pool was sequenced on HiSeq4000 in SE-50bp mode.

### Deep whole genome sequencing

Deep whole-genome sequencing was performed on 56 tumour and matched normal samples, of which 48 passed quality control. Libraries were constructed with ~350-bp insert length using the TruSeq Nano DNA Library prep kit (Illumina) and sequenced on an Illumina HiSeq X Ten System in paired-end 150-bp reads mode. The average depth was 60× (range 40-101 ×) in tumours and 40× (range 24-73×) in matched blood samples.

### Variant calling

sWGS reads were aligned and relative copy-number called as described^38^. Read alignment and variant calling of tagged-amplicon sequencing data were processed as described^38^. Deep WGS samples were processed with bcbio-nextgen^39^ using Ensembl somatic variants called by two methods out of VarDict^40^, Varscan^41^ and FreeBayes^42^. Somatic SNV calls were further filtered based on mapping quality, base quality, position in read, and strand bias as described^43^. In addition, the blacklisted SNVs from the Sanger Cancer Genomics Project pipeline derived from a panel of unmatched normal samples were used for filtering^44^.

### Data download

ICGC: Consensus SNVs and INDELs (October 2016 release), consensus structural variants (v 1.6), consensus copy-number calls (January 2017 release), donor clinical (August 2016 v7-2) and donor histology information (August 2016 v7) for 95 ovarian cancer samples were downloaded from the PCAWG data portal. ACEseq copy-number calls were used for analysis.

TCGA: ABSOLUTE^45^ copy-number profiles from Zack et al^33^ for 402 ovarian cancer TCGA samples were downloaded from Synapse^46^. SNVs for these samples were downloaded from the Broad Institute TCGA Genome Data Analysis Center (Broad Institute TCGA Genome Data Analysis Center: Firehose stddata 2016_01_28 run. doi:10.7908/C11G0KM9, Broad Institute of MIT and Harvard). Donor clinical data were downloaded from the TCGA data portal.

### Absolute copy-number calling from sWGS

After inspection of the *TP53* mutation status and relative copy-number profiles of the 300 sequenced BriTROC-1 samples, 44 were excluded from downstream analysis for the following reasons: low purity (21), mislabelled (7), pathology re-review revealed sample was not HGSOC (3), no detectable *TP53* mutation (13). Of the remaining samples, 57 showed an over segmentation artefact (likely due to fixation). A more strict segmentation was subsequently applied to these samples to yield a usable copy-number profile. Relative copy-number profiles were generated using QDNAseq^47^ and these were combined with mutant allele frequency identified using tagged amplicon sequencing in a probabilistic graphical modelling approach adapted from ABSOLUTE^45^ to generate absolute copy-number profiles. Using Expectation-Maximisation, the model generated a posterior over a range of *TP53* copy-number states, using the TP53 mutant allele frequency to estimate purity for each state. The *TP53* copy-number state which provided the highest likelihood of generating a clonal absolute copy-number profile was used to determine the final absolute copy-number profile. Following absolute copy-number fitting, the samples were rated using a 1-3 star system. 1-star samples (n=54) showed a noisy copy-number profile and were considered likely to have incorrect segments and missing calls. These were excluded from further analysis. 2-star samples (n=52) showed a reasonable copy-number profile with only a small number of miscalled segments. These samples were used (with caution) for some subsequent analyses. 3-star samples (n=147) showed a high-quality copy-number profile that was used in all downstream analyses. The maximum star rating observed per patient was 1-star in 15 patients, 2-star in 26, and 3-star in 91.

### Copy-number signature identification

Preprocessing: The 91 3-star BriTROC-1 absolute copy-number profiles were summarised using six different feature density distributions (Outlined in Figure 1): 1. segment size - the length of each genome segment in 10MB units; 2. Breakpoint number per 10MB - the number of genome breaks appearing in 10MB sliding windows across the genome; 3. change-point copy-number - the absolute difference in CN between adjacent segments across the genome; 4. segment copy-number - the observed absolute copy-number state of each segment; 5. normalised distance of breakpoints from centromere - the distance of a break from the centromere divided by the length of the chromosome arm; 6. length of segments with oscillating copy-number - a traversal of the genome counting the number of contiguous CN segments alternating between two copy-number states. Each of these distributions was then reduced to a set of five variables encoding the shape of the distribution: the first four moments (mean, variance, skewness and kurtosis) and a fifth variable, Hartigan’s dip test^48^, which is a measure of the modality of the distribution. As skew can take a negative value, its absolute value was used. All variables were then scaled to lie between (0,1). This approach resulted in a set of 30 variables representing the copy-number profile of each sample.

Signature identification using NMF. The NMF Package in R^49^ was used to decompose the patient-by-variable matrix into a patient-by-signature and signature-by-variable matrix. A signature search interval of 3-12 was used, running the NMF 1000 times with different random seeds for each signature number. As provided by the NMF Package in R^49^, the cophenetic, dispersion, silhouette, and sparseness coefficients were computed for the signature-by-variable matrix (basis), patient-by-signature matrix (coefficients) and connectivity matrix (consensus, representing patients clustered by their dominant signature across the 1000 runs). 1000 random shuffles of the input matrix were performed to get a null estimate of each of the scores (Supplementary Figure 2). We sought the minimum signature number that yielded stability in the cophenetic, dispersion and silhouette coefficients, and that yielded the maximum sparsity which could be achieved without exceeding that which was observed in the randomly permuted matrices. 8 signatures was deemed optimal under these constraints and was therefore chosen for the remaining analysis.

Assigning signature exposures to samples: For the remaining 26 2-star patient samples, and the 82 secondary patient samples (from patients with 2- or 3-star profiles from additional tumours), the LCD function in the YAPSA package in Bioconductor^50^ was used to assign signature exposures.

### Copy-number signature validation

The variable summary procedure described above was applied to copy-number profiles from two independent datasets: 95 whole-genome sequenced (approximately 40×) HGSOC samples processed as part of ICGC Pan-Cancer Analysis of Whole Genomes Project^21^, and SNParray profiling of HGSOC cases as part of TCGA^33^. The number of signatures was fixed at 8 for matrix decomposition with NMF. A Pearson correlation coefficient was obtained by correlating the signature by feature matrix from each cohort with one derived from the high-quality sWGS copy-number profiles.

### Identification of copy-number signature patterns

The density distributions appearing in Supplementary Figure 7 were used to assist with determining important patterns associated with each signature. These were generated using a weighted kernel density estimator in R where the copy-number features were weighted by their signature exposures for 117 BriTROC-1 cases. We used these, in combination with the variable weights for each signature (Figure 3), to determine which parts of each distribution were important for defining each signature.

### Association of copy-number signature exposures with other features

Of the 48 deep WGS BriTROC-1 samples, 44 had matched 2- and 3-star sWGS copy-number profiles. As the signature exposures from the sWGS were used for analysis, associations were made only with these 44 samples.

#### Mutational signatures

For 44 deep WGS BriTROC-1 samples and 95 ICGC samples, motif matrices were extracted using the SomaticSignatures R package^51^ and the weights of known COSMIC signatures were determined using the deconstructSigs R package^52^. Signatures showing a median exposure of 0 across the samples were removed. The rcorr function in the Hmisc R package^53^ was used to calculate the correlation matrix between the remaining SNV and CN signature exposures.

#### Telomere length

Telomere lengths of 44 deep WGS tumour samples from the BriTROC-1 cohort was estimated using the Telomerecat algorithm^54^. Partial correlation was calculated between telomere length and copy-number signature exposures greater than 10%, with age and tumour purity as covariates, using the ppcor package in R^55^.

#### Tandem duplicator phenotypes

Tandem duplicator phenotype (TDP) scores were calculated for 95 ICGC samples using the method described in Menghi et al^18^. Amplification associated fold-back inversion fraction. For the 95 ICGC samples, the fraction of amplification associated fold-back inversion events per sample was calculated as the proportion of head-to-head inversions within a 100kb window amplified regions (copy number ≥5).

The significance of all observed correlations was estimated from a t-distribution where the null hypothesis was that the true correlation was 0. All reported p-values have been adjusted for multiple testing with Benjamini & Hochberg (BH) method^56^. Comparison plots can be found in Supplementary Figure 5.

### Mutated pathway enrichment analysis

A combined set of 319 samples (39 deep WGS BriTROC-1, 94 ICGC and 186 TCGA) showing at least one driver mutation were used for mutated pathway enrichment analysis. SNVs, INDELs, amplifications (CN>5) or deletions (CN<0.4) affecting 459 ovarian cancer driver genes from IntOGen^57^ were considered bona fide driver mutations if they had TIER1 or TIER2 status based on predictions by Cancer Genome Interpreter^58^ (Supplementary Tables 2 and 3). 137 of the 459 genes were mutated in a least one case. These genes were used to test for enriched pathways in the Reactome database using the ReactomePA^59^ R package with a p-value cutoff of 1 and q-value cutoff of 0.05. Pathways with at least 5 genes were retained. For each pathway, patients were split into two groups: those with mutated genes in the pathways, and those with wild-type genes in the pathways. For each signature, a one-sided t-test was carried out to determine if the signature exposure was significantly higher in mutated cases versus wild-type cases. After multiple testing correction using the Benjamini & Hochberg method (thresholding the p-value <0.1 and the mean difference in exposures >0.06), 113 pathways were significantly enriched (Supplementary Table 4). Manual inspection revealed significant redundancy in the list and 5 representative pathways were selected as a final output (Supplementary Figure 6, Supplementary Table 5).

### Survival analysis

Overall survival in BriTROC-1 patients was calculated from the date of enrolment to the date of death or the last documented clinical assessment, with data cutoff at 1 December 2016. Given that the BriTROC-1 study only enrolled patients with relapsed disease, it was necessary to left truncate the overall survival time. In addition, cases where the patient was not deceased were right censored. Survival data for the ICGC and TCGA cohorts were right censored as required (left truncation was not necessary). The combined samples were split into training (100% BriTROC-1, 60% ICGC and 60% TCGA = 347) and test (40% ICGC and 40% TCGA = 198) cohorts. All of the BriTROC-1 samples were used in the training set to avoid issues calculating prediction performance on left-truncated data. A Cox proportional hazards model was fitted on the training set, with the signature exposures as covariates, stratified by cohort (BriTROC-1, OV-AU, OV-US, TCGA) and age (<51; 52-59; 60-69; >70), using the survival package in Bioconductor^60^. As the signature exposures summed to 1, signature 2 was used to normalise exposures of the remaining signatures.

After fitting, the model was used to predict risk in the test set and performance was assessed using the concordance index calculation in the survcomp package in Bioconductor^42^. An alternative survival analysis approach was also applied, where the maximum signature exposure in each patient was used to assign a patient to either a good prognosis group (signature 1, 7, or 8) or a poor prognosis group (signature 2,3,4,5,6). A second Cox proportional hazards model was fitted on the training set using the groups as covariates to determine a log-rank score. The survfit function was used to generate a Kaplan-Meier plot for the grouping. The survdiff function was used test the performance of this approach in the test set and the survfit function used to generate Kaplan-Meier curves (Figure 4).

### Analysis of copy-number signature changes during treatment

BriTROC-1 samples collected at diagnosis (N=55, prior to treatment) were combined with relapse samples (N=100, at least one line of chemotherapy, Figure 5a). As all diagnosis samples were formalin fixed, and all relapse samples methanol fixed, we first tested if there were differences in signature exposure caused by fixation. Excluding patients with paired samples, Supplementary Figure 10 shows that no significant differences were observed. A two-sided student t-test confirmed this observation for each signature (Supplementary Table 7). We therefore proceeded with two comparisons to highlight differences in signature exposure in response to treatment: 1 - signature exposures compared to number of lines of therapy, and 2 - signature exposures between primary and relapse samples within the same patient. For 1, we fit a linear model with lines of chemotherapy as covariates and signature exposures as output, for resistant and sensitive patients (Figure 5b) using the geom_smooth function from the ggplot2 package in R. A one-sided student t-test was used to determine if there was a difference in means of signature 4 exposures between resistant and sensitive patients at diagnosis (P=0.01, 0 lines of chemotherapy). For 2, we plotted the differences between primary (no therapy) and relapse (chemotherapy) samples for groups of patients with one of each sample (N=36), split by resistant and sensitive cases (Supplementary Figure 8). We performed a two-sided paired t-test to each of these categories (Supplementary Table 6).

### Data and Software Availability

Sequence data that support the findings of this study have been deposited in the European Genome-phenome Archive with the accession code EGAS00001002557. All code required to reproduce the analysis outlined in this manuscript can be found in the following repository: https://bitbucket.org/britroc/cnsignatures.

---

**Supplementary Figure 1.**
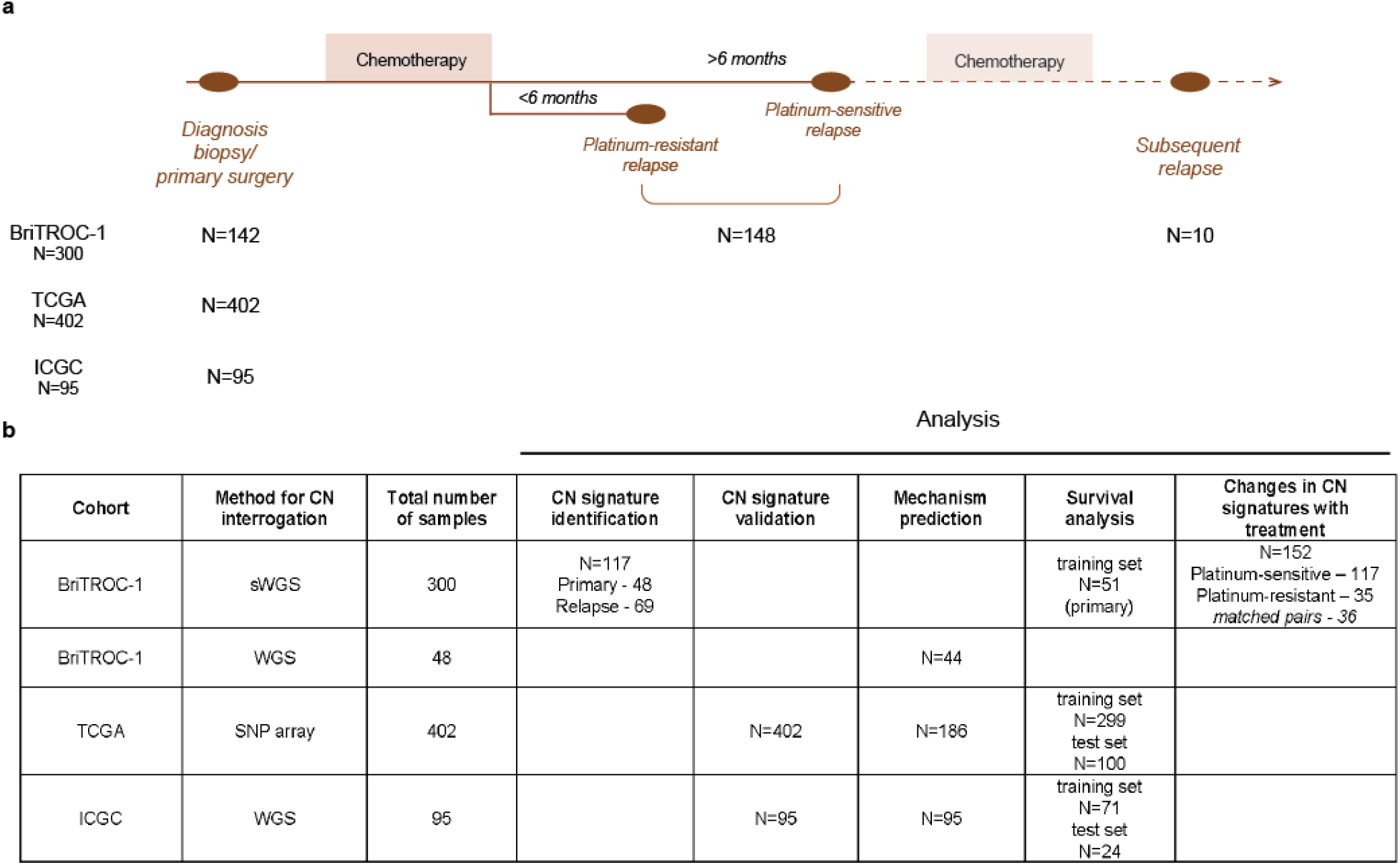
Sample details and workflow. **a** HGSOC datasets used for copy number signature analysis. **b** Samples used for individual analyses. sWGS was performed on 142 BriTROC-1 patients including 300 samples (142 primary, 148 relapse biopsies and 10 biopsies from subsequent progressive disease). 117 patients with high and intermediate quality samples were used for CN signature identification (maximum one sample per patient). CN signatures were initially derived using 91 high quality samples and then applied to the 26 samples with intermediate quality. 48 of the BriTROC-1 relapse samples were also sequenced by deep WGS. Changes in CN signature exposures following treatment were explored in 152 samples from 116 BriTROC-1 patients. (sWGS = shallow whole genome sequencing, WGS = whole genome sequencing, SNParray = single nucleotide polymorphism hybridization array.)

**Supplementary Figure 2.**
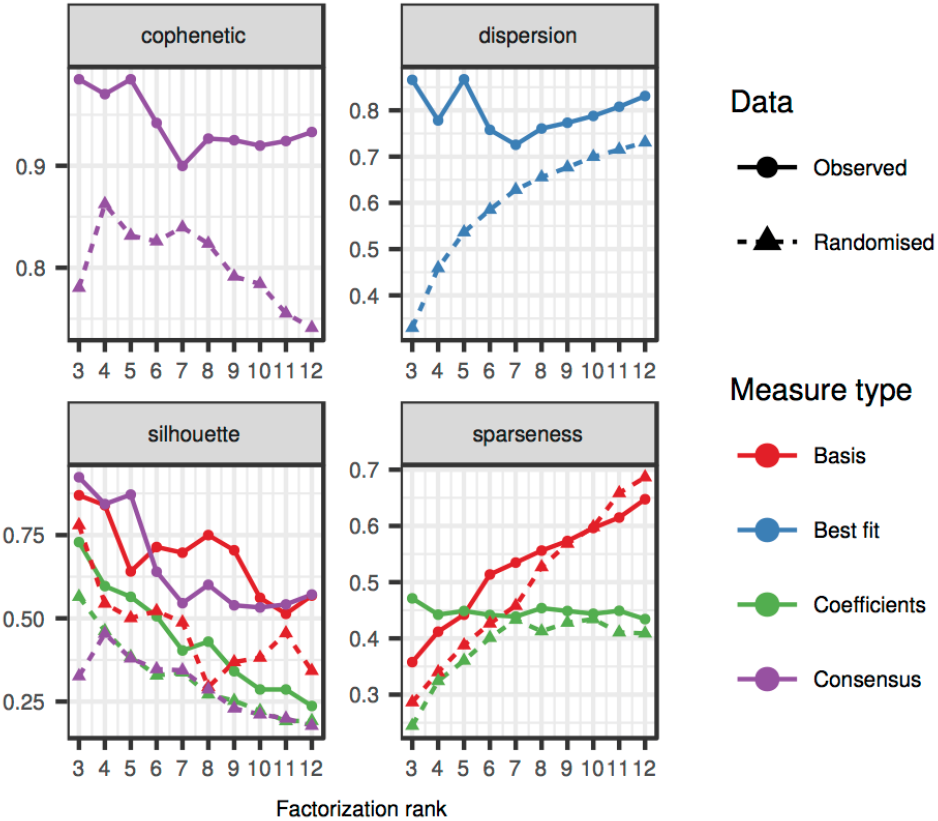
Measures for selecting optimal signature number. A comparison of signature number (x-axis) across four measures for determining optimal signature number. The circle and solid lines represent the results from the BriTROC-1 samples run whereas the triangles and dotted lines represent results from 1000 randomly permuted BriTROC-1 matrices (these can be considered a null measure). Here, basis refers to the signature-by-variable matrix, coefficients refers to patient-by-signature matrix, and consensus refers to the connectivity matrix of patients clustered by their dominant signature across 1000 runs. Best-fit is the run that showed the lowest objective score across the 1000 runs. A value of 8 defines the point of stability in the cophenetic, dispersion and silhouette coefficients, and is the maximum sparsity achievable above the null model.

**Supplementary Figure 3.**
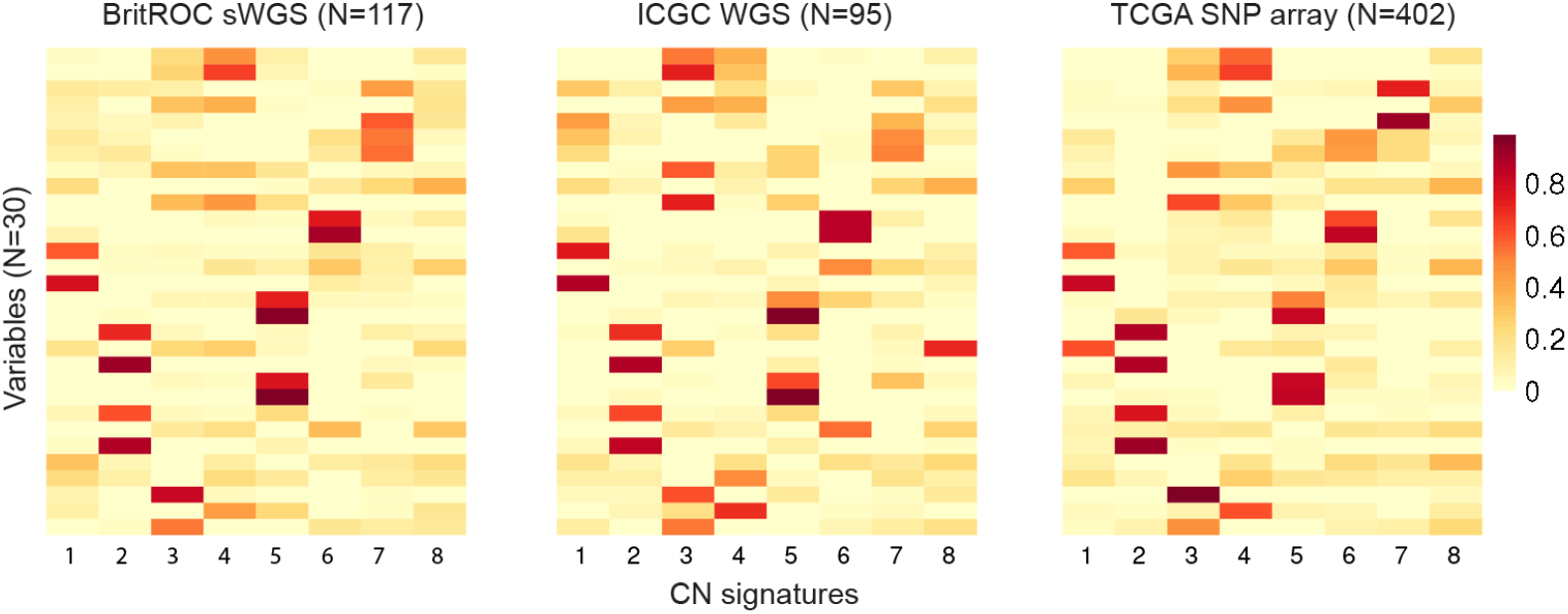
Validation of signature identification. Heat maps show variable weights for copy number signatures in two independent cohorts of HGSOC samples profiled using WGS and SNP array. Correlation coefficients are provided in Supplementary Table 1.

**Supplementary Figure 4.**
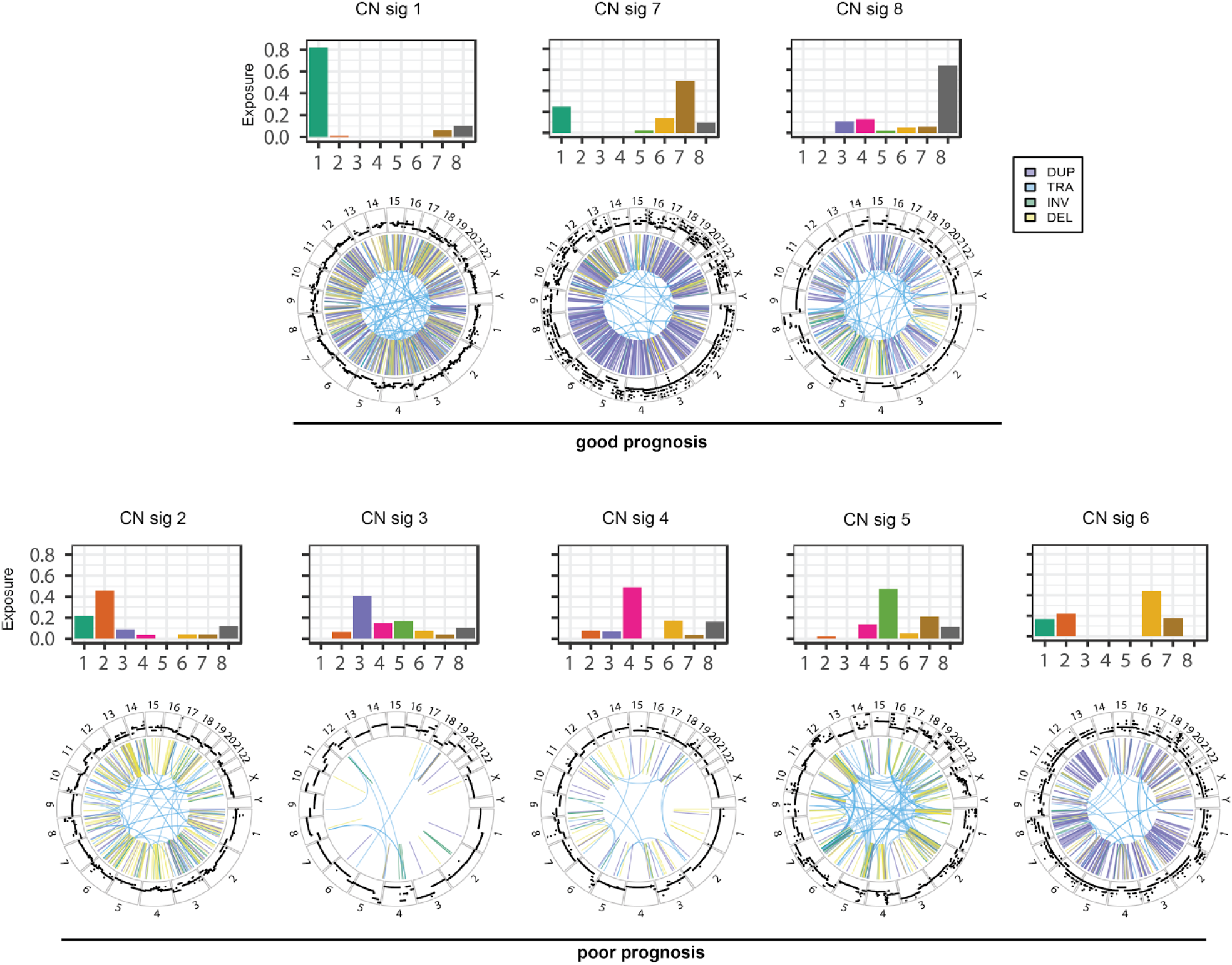
CN signatures profiles and WGS analyses of genomes with dominant CN signatures. Bar plots show the proportion of each CN signature identified in the example case. The outer ring of each circos plot shows copy-number changes. The inner ring shows structural variants: duplications (DUP), translocations (TRA), inversions (INV), and deletions (DEL).

**Supplementary Figure 5.**
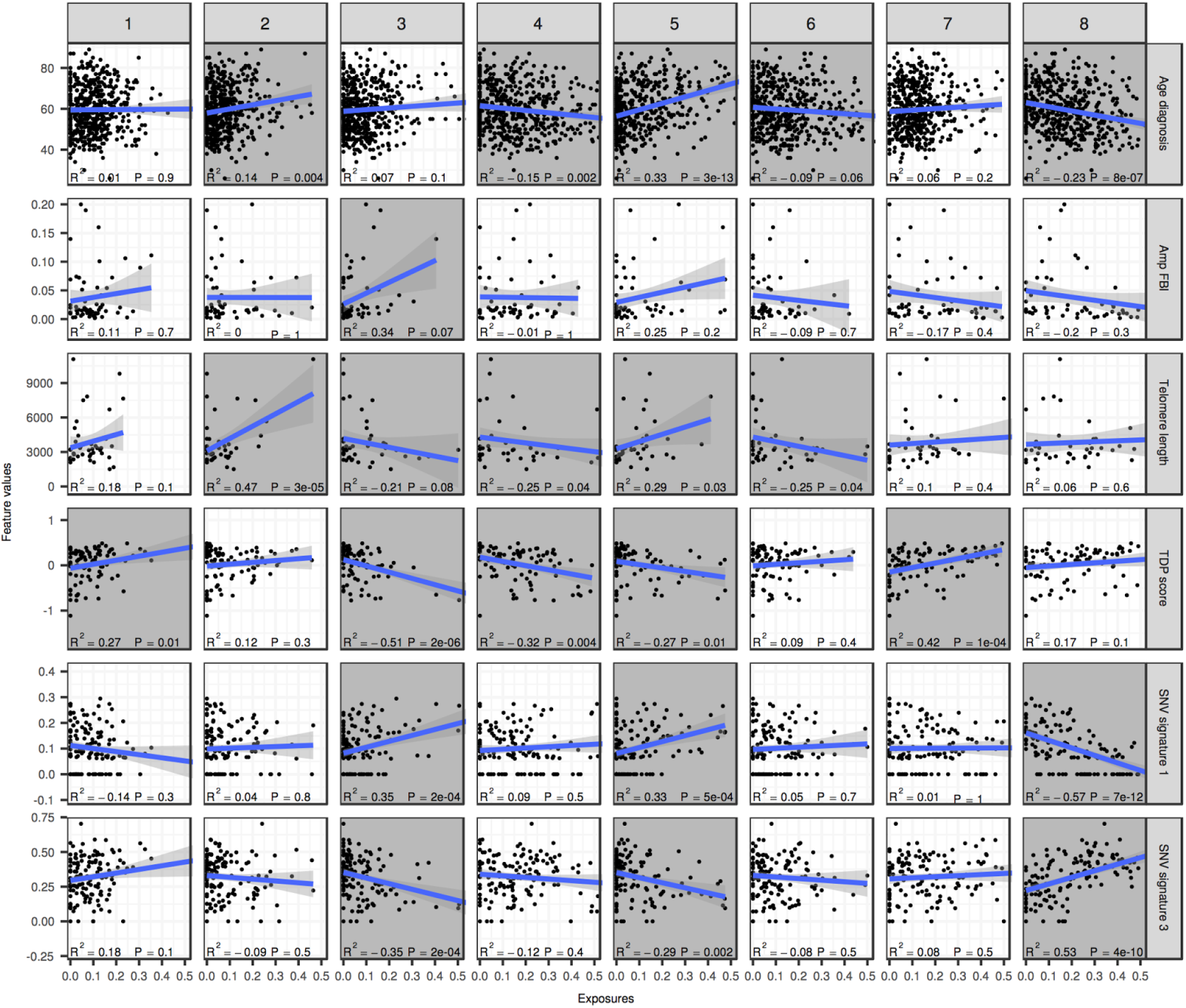
Correlation plots of signature exposures with SNV signatures and other genomic features. Feature values and SNV signatures (right) are correlated with CN signatures exposure (top). Blue lines represent a linear model fit and shading around the lines represent the 95% confidence interval. Shaded panels represent results which are significantly correlated (adjusted P<0.1).

**Supplementary Figure 6.**
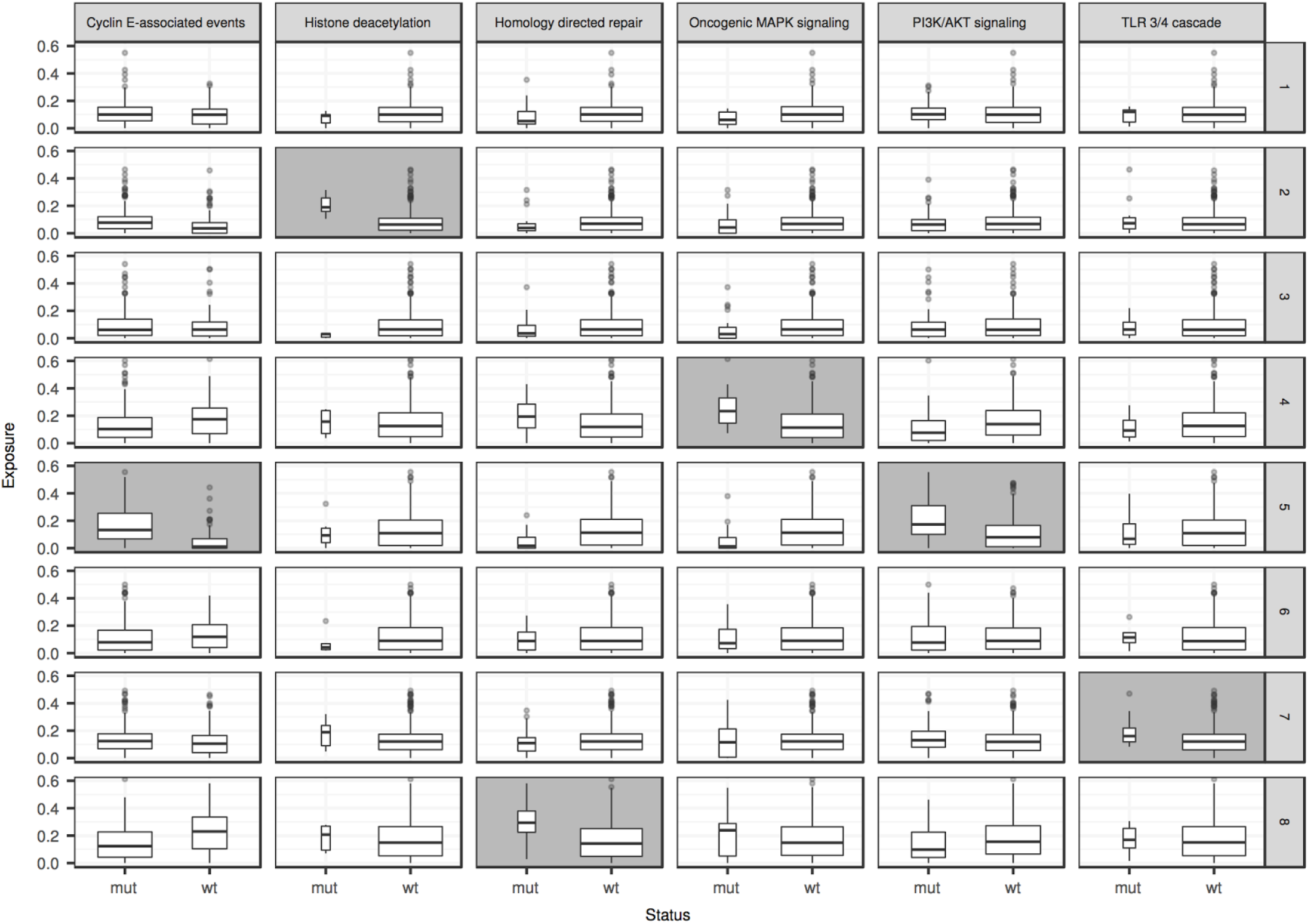
Differences in exposures between cases with mutated pathways versus wild-type. Boxplots representing the signature exposures of cases with mutations (mut) in a given pathway (top) versus those with wild-type alleles (wt). The box widths are proportional to the number of cases (exact numbers can be found in Figure 2). Shaded panels indicate significant differences (adjusted P<0.1, values found in Supplementary Table 4).

**Supplementary Figure 7.**
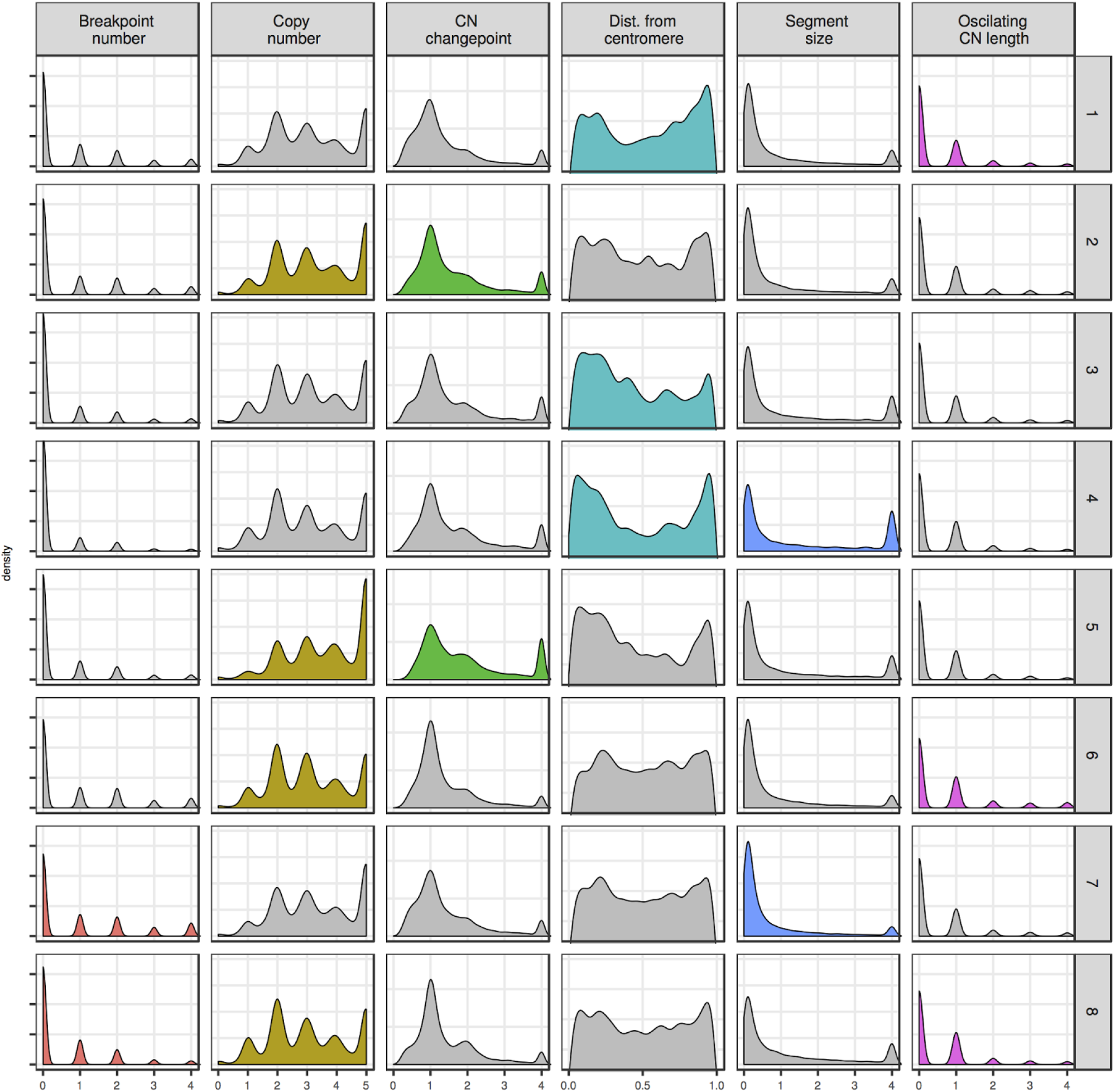
Overview of copynumber feature distributions. Separate density distributions are plotted for each copy-number feature across all eight CN signatures. The distributions that have highly weighted variables (see Figure 3) for each of the feature distributions are coloured. Boxplots represent the signature exposures of paired BriTROC-1 primary tumour samples (P) and relapse samples (R), separated by signature (top) and platinum status at study entry (sensitive vs resistant relapse) (right). Shaded panels indicate significant differences (adjusted P<0.1, values found in Supplementary Table 6).

**Supplementary Figure 8.**
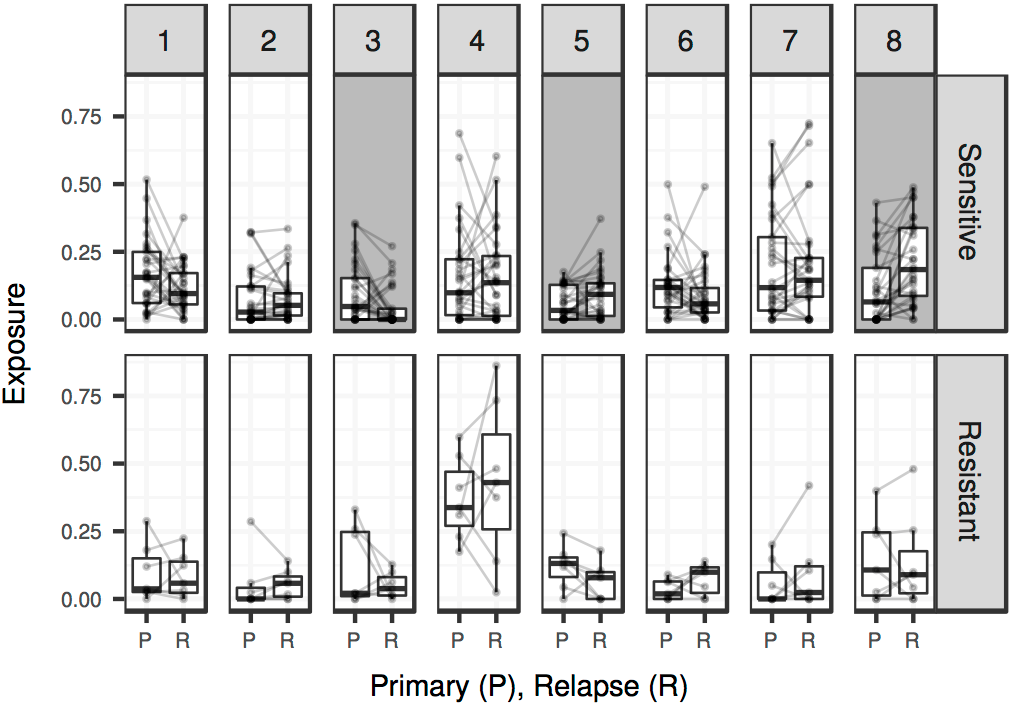
Signature exposure comparison between paired primary and relapsed disease. Boxplots represent the signature exposures of paired BriTROC-1 primary tumour samples (P) and relapse samples (R), separated by signature (top) and platinum status at study entry (sensitive vs resistant relapse) (right). Shaded panels indicate significant differences (adjusted P<0.1, values found in Supplementary Table 6).

**Supplementary Figure 9.**
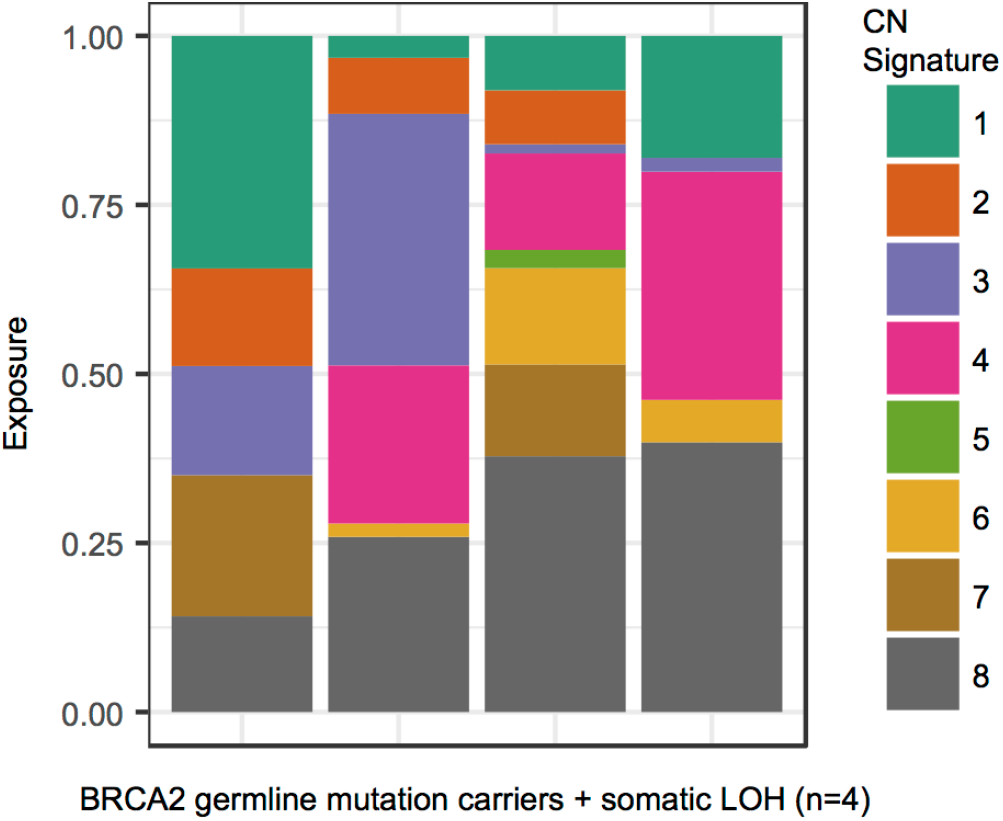
CN signature exposures of 4 BriTROC-1 patients with germline *BRCA2* mutations and somatic loss of heterozygosity. Stacked bar plots show copy-number signature exposures for four BriTROC-1 cases with confirmed pathogenic germline *BRCA2* mutations and somatic loss of heterozygosity (LOH).

**Supplementary Figure 10.**
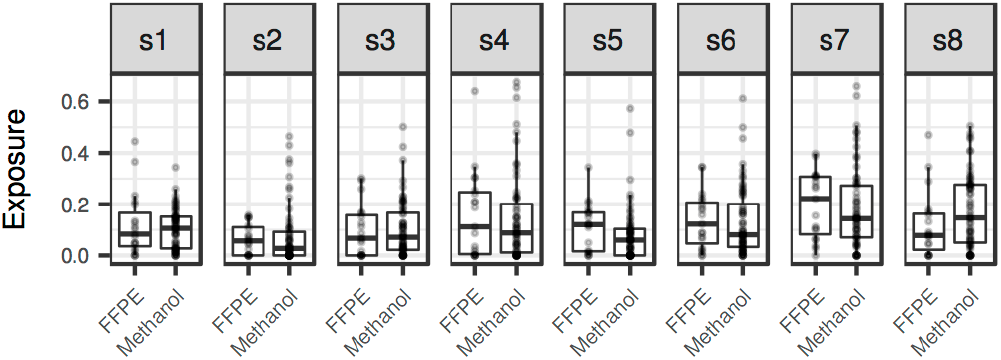
Signature exposure comparison between FFPE and methanol fixed samples. Boxplots represent the signature exposures of primary FFPE samples and relapse methanol fixed samples, separated by signature (top). There are no significant differences observed due to fixation differences (adjusted P>0.1, values found in Supplementary Table 7).

